# Small-molecule inhibitors targeting prokaryotic ClpQ protease in *Plasmodium* display anti-malarial efficacies against blood and liver stages

**DOI:** 10.1101/2022.05.12.491652

**Authors:** Shaifali Jain, Mohd Asad, Reto Caldelari, Gaurav Datta, Shweta Singh, Sumit Rathore, Rebecca R. Stanway, Volker T. Heussler, Asif Mohmmed

**Affiliations:** International Centre for Genetic Engineering and Biotechnology, New Delhi 110 067, India; Institute of Cell Biology, University of Bern, Bern, Switzerland; All India Institute of Medical Sciences, New Delhi, India

**Keywords:** Malaria, Drug-target, mitochondrion, ClpQ protease, Malaria Box

## Abstract

The ATP-dependent ClpQY system is a prokaryotic proteasome-like machinery in the mitochondrion of malaria parasite, *Plasmodium*. This protease system is identified as a validated target for malaria intervention. Here we have identified drug-like small chemical compounds, pyrimidotriazine derivatives, as specific inhibitors of *P. falciparum* ClpQ (*Pf*ClpQ) having effective parasiticidal efficacies on asexual and liver stages. Screening of the ‘Malaria Box’ compound library, using a robust *in vitro* activity assay, identified a hit compound targeting the *Pf*ClpQ protease with IC_50_ at ~750nM. Further, Structure-Activity Relationship (SAR) analysis, using a small library of derivatives of the hit compound, identified three potential lead compounds, ICGEB-L1, ICGEB-L2 and ICGEB-L3, which inhibited enzyme activity with IC_50_ in 100-200nM ranges. The three lead compounds effectively inhibited *in vitro* asexual stage parasite growth in similar concentration ranges with ~100-fold selectivity over mammalian cell line; in addition, these compounds effectively inhibited *in vitro* growth and maturation of hepatocytic stage parasites with EC_50_ in low nano-molar ranges. These compounds were found to effectively block the development of trophozoite stages to schizont stage, concurrently arresting development and segregation of parasite mitochondria, in asexual as well as hepatocytic stages of the parasite. These morphological and developmental effects of the drugs on parasites mimic the data from genetic ablation of *Pf*ClpQ. Importantly, all three compounds showed significant *in vivo* anti-parasitic efficacies on blood stages of *P. berghei* mouse malaria model. Overall these results suggest that these *Pf*ClpQ targeting compounds are promising candidates to develop new anti-malarials.

## Introduction

Malaria remains to be one of the major infectious diseases, especially affecting children in tropical and subtropical regions of the world. Malaria caused about 241 million cases globally resulting in 627,000 deaths (1–3). Although, artemisinin based combination therapies resulted in major decline in malaria cases in last few years, there is rapid emergence of parasite strains resistant to most of commonly used antimalarials including artemisinin combination therapies. Therefore, there is an urgent need to identify new lead drug molecules and develop new antimalarials. Selection of unique and essential metabolic pathways in the parasite is a prerequisite to develop unique and specific drug targeting the parasite. Metabolic pathways in two parasite organelles of prokaryotic origin, the apicoplast and the mitochondrion, are considered to be important drug targets to obstruct the parasite development and segregation (4, 5).

ATP dependent protease machineries include eukaryotic 26S proteasome system, which are large multi-subunit protein complexes (6, 7). In eukaryotic cells, the 26S proteasome consists of a protease component, 20S proteasome, and the 19S proteasome component which is chaperon like ATPase. The 20S complex consists of four stalked heptameric rings of α- and β-type subunits that form a barrel shape structure. The β-type subunits contain the proteolytic active sites and form the inner rings of the complex. The 19S complex contains 17 subunits, forms a hetero-hexameric ring at either end of the 20S protease barrel. The 19S complex recognizes polyubiquinated proteins and unfolds and translocates them into the 20S particle for degradation (6) The 26S proteasome machinery plays essential roles in controlling the levels of key regulatory proteins and in the elimination of abnormal polypeptides in eukaryotic cells. Similar multi-subunit protein complexes in archaea and eubacteria carry out these tasks, these machineries include ClpAP, ClpXP, and ClpQY (HslVU) proteases (7). The ClpQY machinery, a multimeric ATP dependent protease system, in the prokaryotes, consists of two stacked hexameric rings of ClpQ protease with hexameric ring of AAA type ATPase, ClpY, which form cap on one or both sides of the protease barrel (8). The ATPases caps act as chaperons to unfold the substrate proteins for their entry into the protease barrel to get degraded. Our group has earlier carried out detailed studies to understand the biochemical and functional role of the ClpQY machinery in the parasite (ref).

*P. falciparum* ClpQ, *Pf*ClpQ, is a threonine protease localized in the parasite mitochondria (9, 10); detailed genetic and chemical ablation studies have shown its essentiality for parasite survival and hence validated it to be a drug-target (9, 11). In the present study, we have identified drug-like small chemical compounds targeting PfClpQ, which have effective parasiticidal efficacies. We show that inhibition of activity of ClpQ protease in malaria parasite using these small chemical compounds results in mitochondrial developmental arrest as well as death/growth inhibition of asexual blood stage and liver stage parasites, and therefore these compounds can be utilized to develop antimalarial therapeutics.

## Results

### Development of robust *in vitro* activity assay of *Pf*ClpQ protease, screening of the ‘Malaria Box’ and identification of a hit compound

To determine the enzymatic activity of *Pf*ClpQ, we expressed a recombinant protein corresponding to its protease domain (41aa-338aa) as described earlier (9). The corresponding recombinant protein was purified and separated on SDS-PAGE. On the gel it migrated at the predicted size of ~20kDa (Fig. 1A). The purified recombinant *Pf*ClpQ was used to standardize a fluorescent-based activity assay to quantify its protease activity. A fluorescent peptide substrate-based activity assay was standardized using a tripeptide threonine protease substrate which bears covalently linked fluorophore and quencher on either end, N^α^-Benzyloxycarbonyl-glycine-glycine-leucine-7-amido-4-methylcoumarin (Z-GGL-AMC). The enzyme activity is recorded as the increase in fluorescence of the reaction due to proteolytic cleavage of the peptide to release the fluorophore.

**Figure 1:**
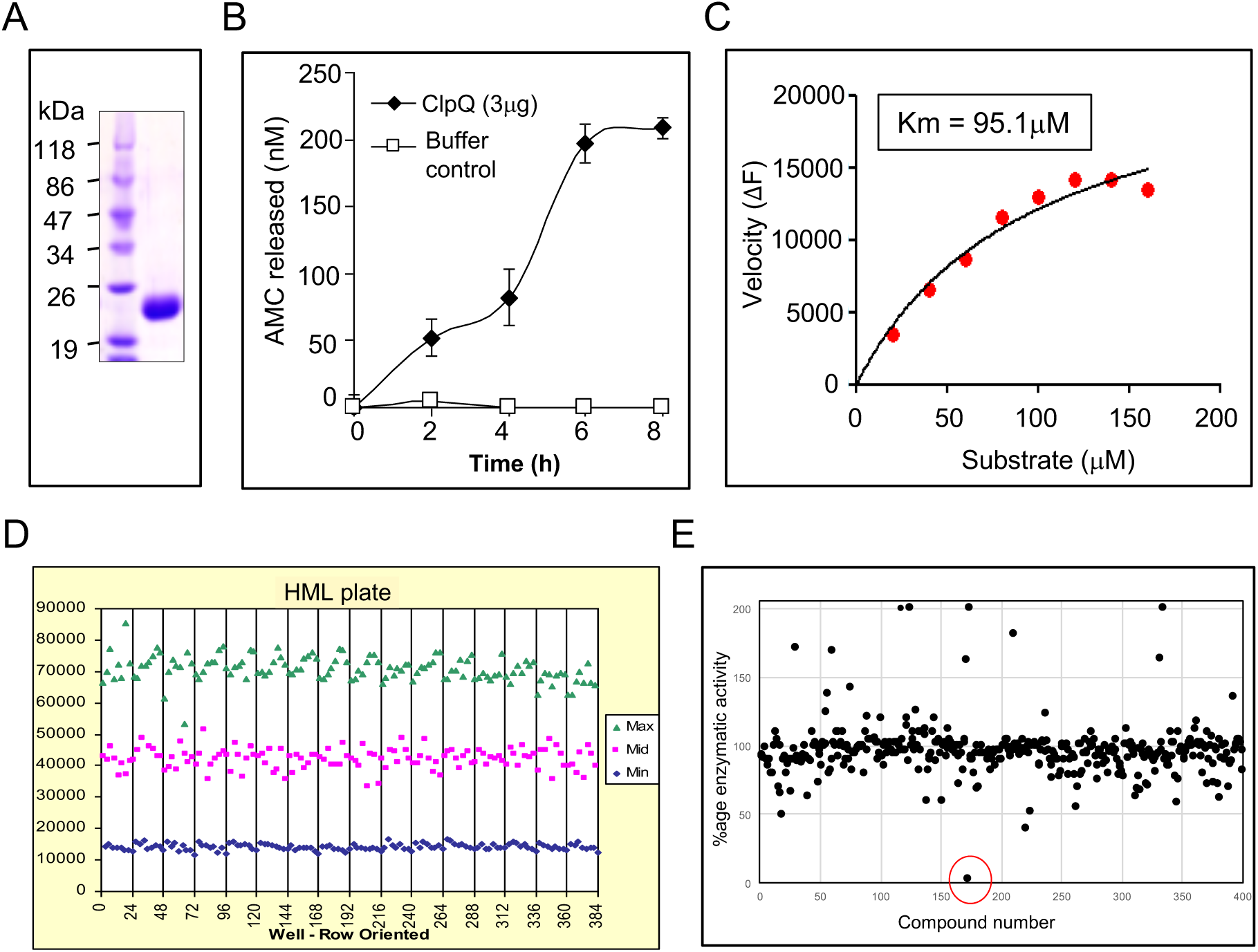
Protease activity assay of recombinant *Pf*ClpQ enzyme and screening of MMV Malaria Box library. (A) Coomassie stained SDS-PAGE showing purified recombinant *Pf*ClpQ. (B) Graphical representation of protease activity of PfClpQ as assessed by cleavage of fluorogenic peptide substrates, Suc-Gly-Gly-Leu-AMC; the activity was determined by estimating release of 7-amido-4-methylcoumarin (AMC) in time dependent manner. (C) Line graph of Michelis-Menten fit for *Pf*ClpQ activity using different concentrations of the substrate; the *K*m value of *Pf*ClpQ was found to be 95.1µM. (D) Robustness of the activity assay was assessed by estimating Z-value carried out in HML format in two 96-well plates for two consecutive days; fluoresce units for the activity assay of each well of the four plates are shown. The Z-value was found to be ~0.8, suggesting robustness of the assay. (E) Primary screening of Malaria Box compound library using *in vitro* activity assay of *Pf*ClpQ. The screening was carried out at 10µM concentration of each compound; and percentage enzymatic activity for each compound as compared to control set is presented in the dot plot. One of the compounds, MMV306025 was identified to inhibit >90% enzyme activity, which are labelled as CN172.

To standardize the optimum activity assay conditions, various buffers conditions encompassing a range of pH and reductant concentrations in the reaction, were used. The protease showed maximal activity at pH 8.0 in HEPES-CaCl_2_-MgCl_2_ buffer with 3mM DTT. These assay conditions were then followed for all activity-assay based experiments. The recombinant *Pf*ClpQ showed concentration dependent activity in the assay, confirming the predicted threonine protease activity of *Pf*ClpQ (Fig. 1B). The *K*m for the threonine protease activity with Z-GGL-AMC substrate was estimated to be ~95µM. The robustness of this activity assay was assessed, so that it can be used for high throughput screening (Fig. 1C). For this, 96-well plate assays were conducted for a period of three days and the Z-value calculated based on the readouts from duplicate plate assays on all three days, which was found to be 0.8, clearly indicating the robustness of the assay (Fig. 1D) thus also suggesting it could be used for high throughput screening of compounds.

The antimalarial compound library “Malaria Box” obtained from Medicines for Malaria Venture (MMV, Switzerland) consists of 200 drug-like and 200 probe-like compounds. The library was screened for identification of potential inhibitors over the PfClpQ protease activity assay at final concentrations of 10μM (Fig. 1E). Out of these four hundred compounds, one of the compound-172 (MMV306025; labelled as CN172; methyl 4-{1-ethyl-6-methyl-5,7-dioxopyrimido[5,4-e][1,2,4]triazin-3-yl}benzoate) showed maximal inhibition of *Pf*ClpQ protease activity at 10μM concentration, i.e., ~97%; this compound was selected for further studies. The IC_50_ for CN172 was calculated to be 739nM (Fig. 2A) and its *K*i was found to be 440nM.

**Figure 2:**
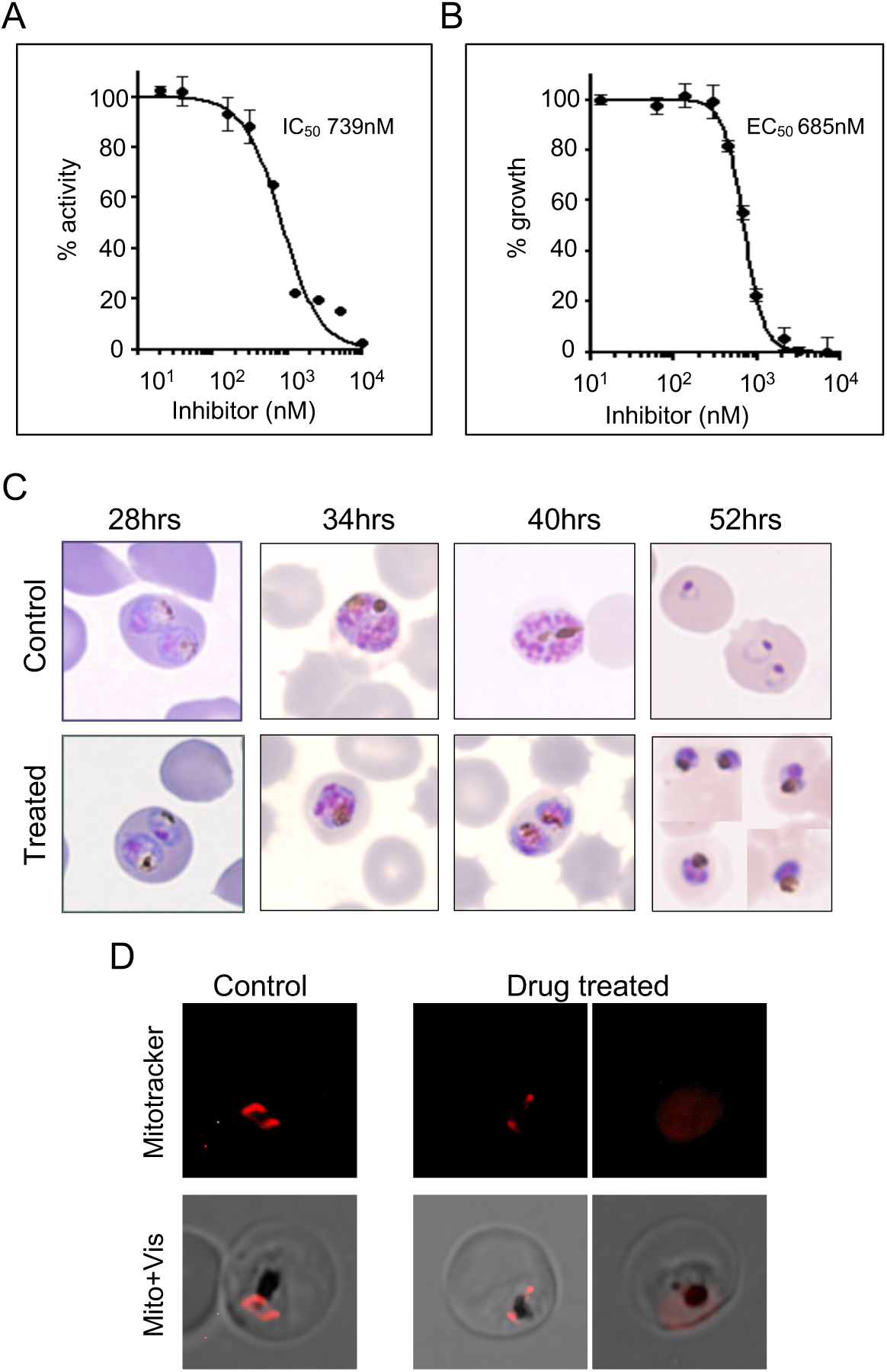
Malaria box compound MMV306025 (CN172) is idenfied as a hit compound targeting ClpQ protease. (A) Concentration dependent inhibition of ClpQ protease activity by hit compound CN172, identified in primary screening of Malaria Box library. (B) Concentration dependent inhibition of asexual blood stages *P. falciparum* parasite growth by compound CN172. (C) Microscopic images of Giemsa-stained parasites showing effect on morphology at different time points after invasion, for parasite culture grown in presence of CN172 (treated) or with solvent alone (control). (D) Fluorescence microscopic images of parasites grown in presence of CN172 (treated) or with solvent alone (control) and stained with MitoTracker. Treated parasites showed poorly stained and punctuate mitochondria as compared to control showing elongated mitochondria.

The “Malaria Box” compounds are known to show parasiticidal efficacies with EC_50_ at low μM range; we assessed the EC_50_ of CN172 compound which was found to be 685nM (Fig. 2B). Further, to study the effect these compounds on parasite development and morphology, parasite in continuous treatment (8-54hpi) set were assessed for developmental-stage profile during the asexual stage cycle. In solvent control set, parasites developed from ring to trophozoite to mature schizont, subsequently merozoites released from these schizonts invaded fresh RBCs and developed into ring stage parasites, which led to about 7 times increase in the total parasitemia. In the drug treated sets most of the ring stage parasites were able to develop into trophozoite stage parasites as in the solvent control sets; however, >50% of these trophozoites were not able to develop into mature schizonts (Fig. 2C), these parasites were seen as stressed trophozoites or abnormal unruptured schizonts. Furthermore, treatment with this compound hindered mitochondrion development in blood stage parasites (Fig. 2D).

### SAR and selection of potential lead compounds/improved hits

The core structure based upon the hit compound CN172 is a pyrimidotriazine; three side chains are identified in the core structure which can be utilized for derivatization of the hit compound. A set of 28 compounds with variations in one of more of these side chains of CN172, were identified from MolPort database and procured. These compounds were screened using the standardized *in vitro* activity assay for *Pf*ClpQ protease. A total of 12 compounds were identified from these results which showed concentration dependent inhibition of enzyme activity with IC_50_ nearly equal or lower than the hit compound CN172. Further, these compounds were assessed in secondary assays to inhibit parasite growth *in vitro* using range of concentrations. Three compounds : MolPort-002-323-496 (1,6-dimethyl-3-(pyridin-4-yl)-1H,5H,6H,7H-pyrimido[5,4-e][1,2,4]triazine-5,7-dione); MolPort-002-323-499 (3-(4-chlorophenyl)-1-ethyl-6-methyl-1H,5H,6H,7H-pyrimido [5,4-e] [1,2,4] triazine-5,7-dione); and MolPort-002-338-765 (1-ethyl-6-methyl-5,7-dioxo-3-[(1E)-2-phenylethenyl]-1H,5H,6H,7H-pyrimido[5,4-e][1,2,4]triazin-4-ium-4-olate) were identified that showed EC_50_ in parasite growth inhibition assay lower than parent compound and within 100-200nM range, which was similar to their respective IC_50_ in activity assays (Fig. 3A, 3B; Table 1). All three compounds were also screened for their selectivity on parasite cells as compared to mammalian cell lines; mammalian cells tolerated these compounds at ≥100-fold higher EC_50_ values; Table 1). These three compounds were further analyzed and for simplicity labelled as ICGEB-L1, ICGEB-L2 and ICGEB-L3.

**Table 1:** *Pf*ClpQ inhibitors identified and their efficacies. The IC_50_ and EC_50_ values of selected compounds, on ClpQ enzymatic activity and growth of asexual blood stage *P. falciparum* respectively, are given. EC_50_ of each compound on mammalian cell line is also indicated.

| Compound ID | IC<sub>50</sub> (nM) to inhibit enzymatic activity of *Pf*ClpQ* | EC<sub>50</sub> (nM) to inhibit *in vitro* growth of *P. falciparum* * | EC<sub>50</sub> (nM) to inhibit *in vitro* growth of HeLa cells* |
| --- | --- | --- | --- |
| MMV306025 (CN172) | 739.5 (6.037) | 685.4 (2.54) | 9577 (30.23) |
| ICGEB-L1 | 76.85 (2.637) | 70.59 (3.12) | 8396 (29.34) |
| ICGEB-L2 | 184.3 (10.48) | 116.4 (19.26) | 9933 (42.78) |
| ICGEB-L3 | 158.7 (10.03) | 193.7 (3.02) | 10016 (12.23) |

\* values in bracket indicate SD

**Figure 3:**
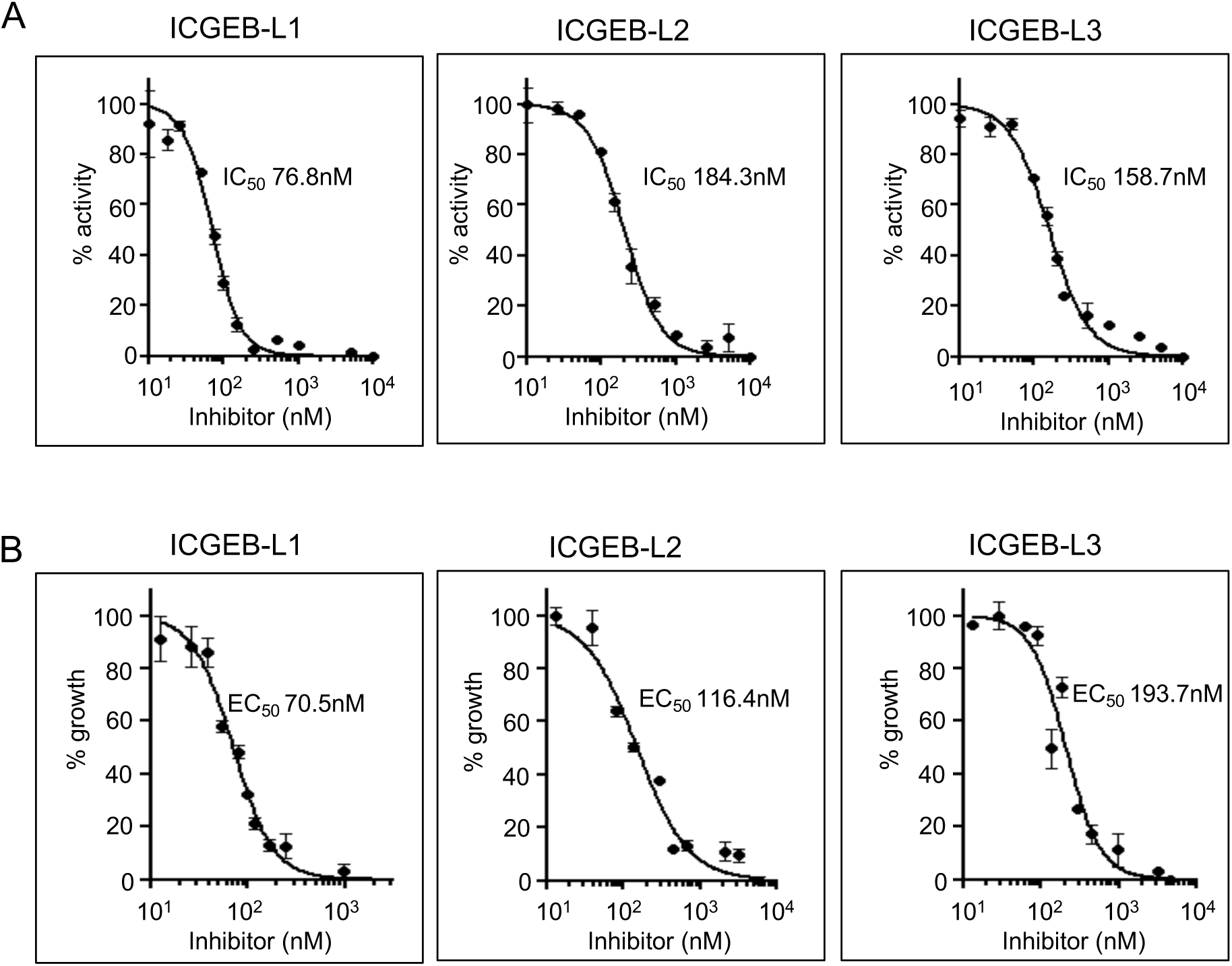
Selected hit compounds, ICGEB-L1, ICGEB-L2 and ICGEB-L3, inhibit *Pf*ClpQ protease activity as well as parasite growth. (A) Graph showing concentration dependent inhibition of *Pf*ClpQ protease activity by ICGEB-L1, ICGEB-L2 and ICGEB-L3 compounds. (B) Graph showing concentration dependent inhibition of asexual blood stages *P. falciparum* parasite growth by ICGEB-L1, ICGEB-L2 and ICGEB-L3 compounds.

### Stage specificity of effect of selected compounds on blood stage parasites

To assess the stage dependent parasiticidal effect of selected compounds on blood stage parasites, highly synchronized *P. falciparum* parasites (8hpi) were treated with each of the compound at concentration ~EC_50_ values for different time-periods of intraerythrocytic life cycle (Fig. 4). The parasite growth inhibition was compared with the treatment with respective compound at concentration ~EC_50_ values for the complete cycle. Treatment with each of the three compounds for 20-32hpi as well as 32-44hpi caused growth inhibition similar to the continuous treatment (8-54hpi) for the respective compound (Fig. 4); however, when the parasites were treated for period of 8-20hpi, there was only 10-20% growth inhibition. These results suggest that treatment with these compounds at trophozoite and schizont stage parasites cause irreversible growth inhibition of the parasites. Overall these results suggest stage specific effect of the ClpQ targeting compounds.

**Figure 4:**
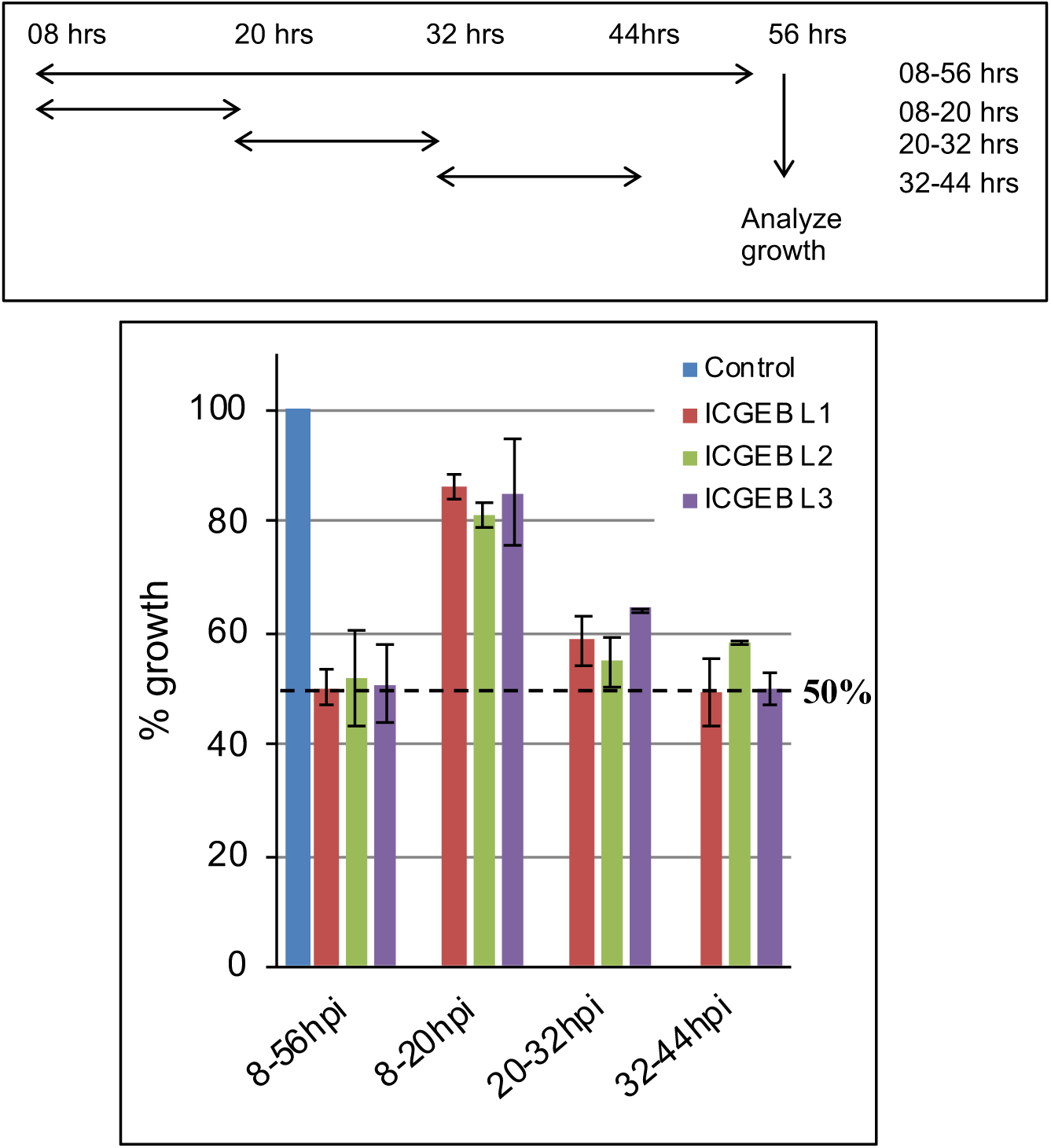
Stage specific growth inhibition of asexual blood stage parasites by selected hit compounds (ICGEB-L1, ICGEB-L2 and ICGEB-L3). Parasite were treated for each of the compound at concentration ~IC_50_ for different time periods during the parasite cell cycle and growth was determined as new ring stage parasite, percentage growth is presented as compared to parasite set treated with solvent control. Schematic showing different duration of treatments of drugs to the EEF is shown.

We also assessed effect of selected compounds on growth of *P. berghei* blood stage parasites *in vitro* using a schizont development assays. The parasites in control wells were able to develop into mature trophozoite/schizont stages. However, all three compounds inhibited *P. berghei* blood stage parasite development in similar manner as in the case of *P. falciparum*; the parasite development from trophozoite to schizont was severely affected in treated parasites sets. The three compounds showed EC_50_ for inhibition of parasite development at concentrations less than half of their respective *Pf*-IC_50_ values.

#### Intra-hepatocyte parasite growth and development is inhibited by selected compounds

ICGEB-L1, ICGEB-L2 and ICGEB-L3 were assessed for their anti-parasitic efficacies on intra-hepatocytic stages; potency of each drug to inhibit development of exoerythrocytic forms (EEF) was analysed using *in vitro P. berghei* sporozoite (Pb-mCherry_*hsp70*_, parasites constitutively expressing mCherry; 12) infection of HepG2 cells. All three compounds showed inhibition of EEF development already at 24h (Fig. 5A) and more pronounced at 48h (Fig. 5B, C) at these concentrations as evident mean infected cell sizes; ICGEB-L3 was found to be most effective in reducing parasite growth over 100μm^2^ at its ~Pf-EC_50_ dose (170nM). ICGEB-L1 and ICGEB-L2 showed a significant growth inhibition at 80nM and 110nM (~Pf-EC_50_) respectively; however, these compounds were not as efficient as ICGEB-L3. However, with increased concentrations (3× Pf-EC_50_) these drugs also efficiently inhibited growth. Parasites also showed a dose dependent reduction in numbers at 48h for all drugs (Fig. 5B). Having assessed the concentration-dependent effect, the effect of different treatments (each compound at concentration of Pf-EC_50_, 3× lower and 10× lower) was examined with the output being completion of liver stage development, as determined by detached cell formation at 65hpi. In this assay, all the three compounds showed >80% reduction in parasite development at Pf-EC_50_ concentration (100-200nM); in addition, compounds ICGEB-L1 and ICGEB-L3 showed >50% inhibition even at one-third of these concentrations (Fig. 6A).

**Figure 5:**
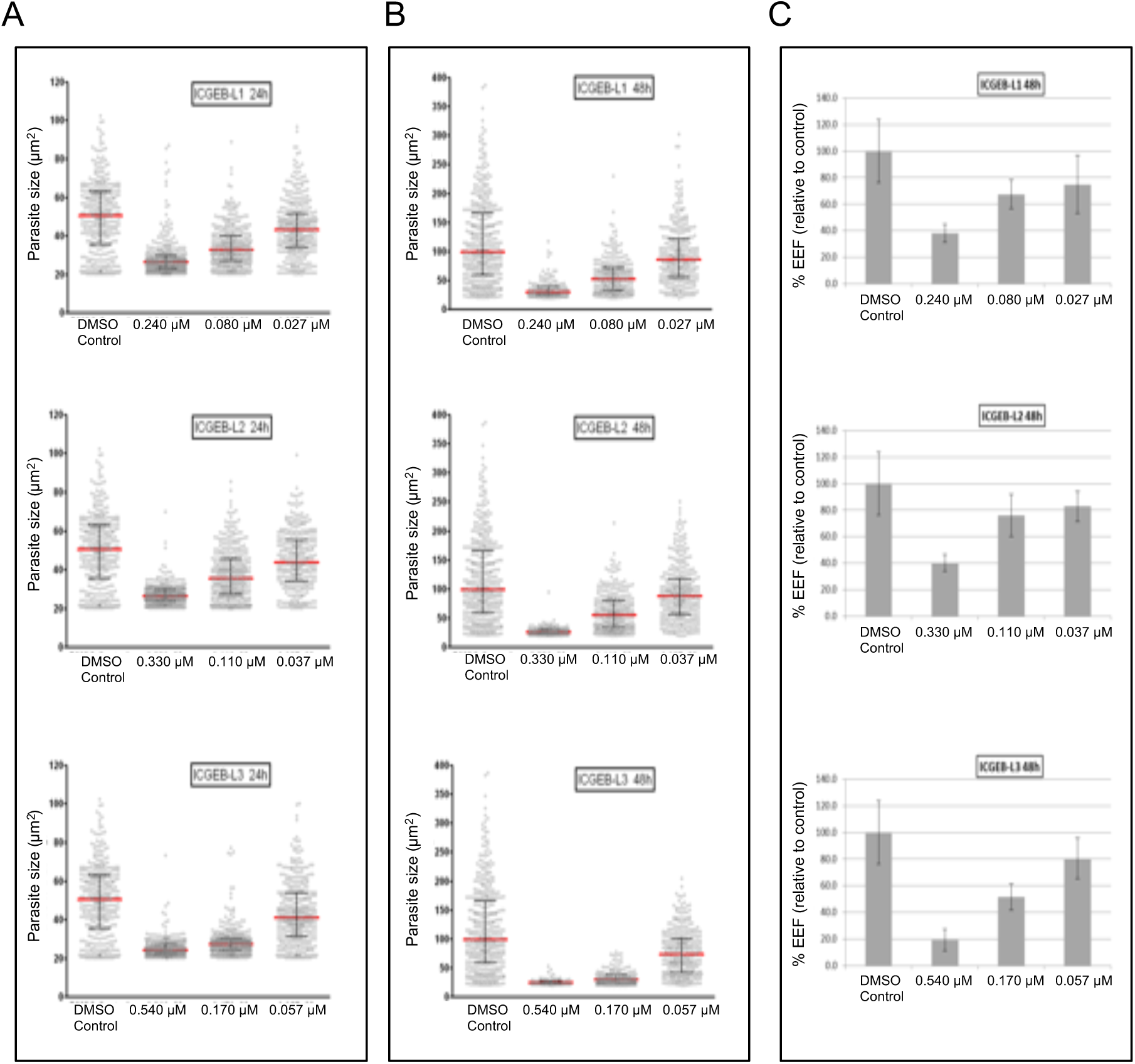
Inhibition of intra-hepatocyte parasite growth by selected compounds. HepG2 cells were infected with *P. berghei* sporozoites and grown in presence of different concentrations of selected compounds; the EEF sizes were determined at 24h after infection (A), and 48h after infection (B); dot plot of ‘In Cell Analyzer’ data of EEF sizes is shown, mean EEF size is marked. (C) Total EEF numbers counted at 48h in each set, shown relative to control.

**Figure 6:**
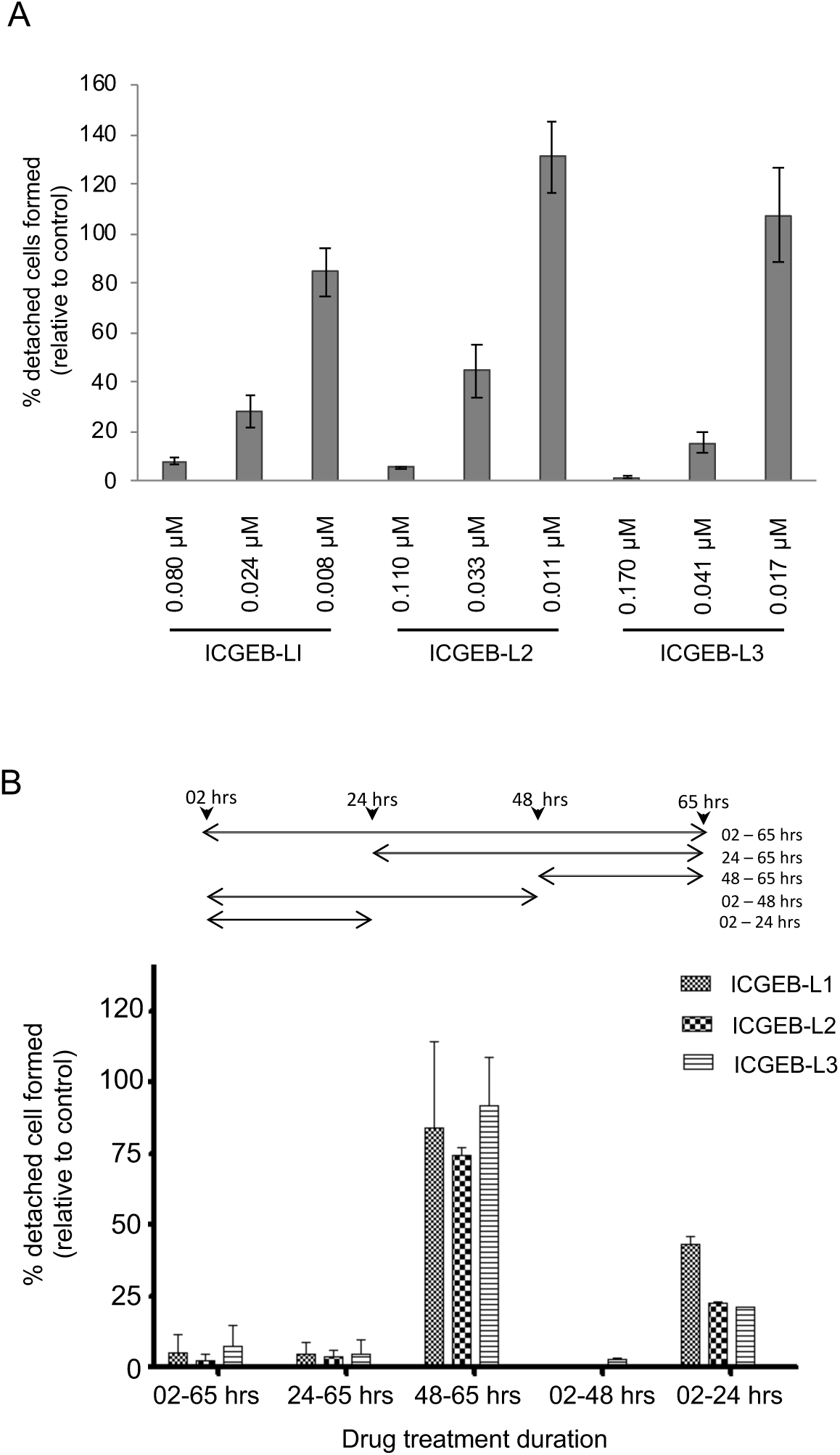
Effect of selected compounds on completion of *in vitro* growth and development of intra-hepatocytic parasites. (A) Inhibition of intra-hepatocyte parasite development by selected lead compounds. Parasite growth was determined by detached cell formation and expressed as compared to control. (B) Inhibition of intra-hepatocyte parasite development by treatment with selected lead compounds for different developmental stages of intra-hepatocyte parasites. Schematic showing different duration of treatments of drugs to the EEF is also shown. Percentage of the intraheptocytic parasites determined by detached cell formation, as compared to control.

#### Inhibition of different developmental stages of intra-hepatocyte parasites

To study the effect of the compounds on different developmental stages of intra-hepatocyte parasites, the compounds were added for varying time periods after sporozoite infection of HepG2 cells (Fig. 6B) with the output being completion of liver stage development (detached cell analysis); each of the compound was used at respective Pf-EC_50_ concentrations (80nM for ICGEB-L1, 110nM for ICGEB-L2, and 170nM for ICGEB-L3) and the treatment windows selected approximately corresponding to trophozoite (6-24hpi), schizont (24-48hpi) and mature schizonts/completely mature detached cells (48-65hpi). As shown above, continuous treatment of cells (2-65h) with any of the three compounds caused 80-90% reduction in parasite development; similar results were obtained when the drugs treatment was during schizont development stages (2-48hpi). However, the effects were less pronounced for drug treatment given during trophozoite developmental stages (2-24hpi), in these sets the ICGEB-L2 and ICGEB-L3 showed ~75% growth inhibition, whereas ICGEB-L1 caused ~55% growth inhibition. Almost no effect was seen when the cells were treated after development of mature stages of the parasites (48-65hpi) (Fig. 6B). These results suggest that the drugs act mainly on the development of schizont stage of the intra-hepatocyte parasites.

#### Effect of selected compounds on mitochondria development during intraerythrocytic and intrahepatocytic stages

Since *Pf*ClpQ is known to play a key role in mitochondrial growth and division (11), we assessed any effect of these ClpQ targeting compounds on parasite mitochondria. *P. falciparum* parasite line expressing HSP60-GFP fusion protein (13) are used for these experiments. Tightly synchronized parasites at ring stages (6-8hpi) were treated with each of the compound and the parasites were visualized after 24h of growth (trophozoite stages). In the solvent control set, the mitochondria showed normal growth and development; in majority of these parasites at trophozoite stage (24 h after treatment) the mitochondria appeared as elongated structures. However, in treated sets the mitochondrial development was severely affected and parasites harbored under-developed or abnormal mitochondria (Fig. 7A, B).

**Figure 7:**
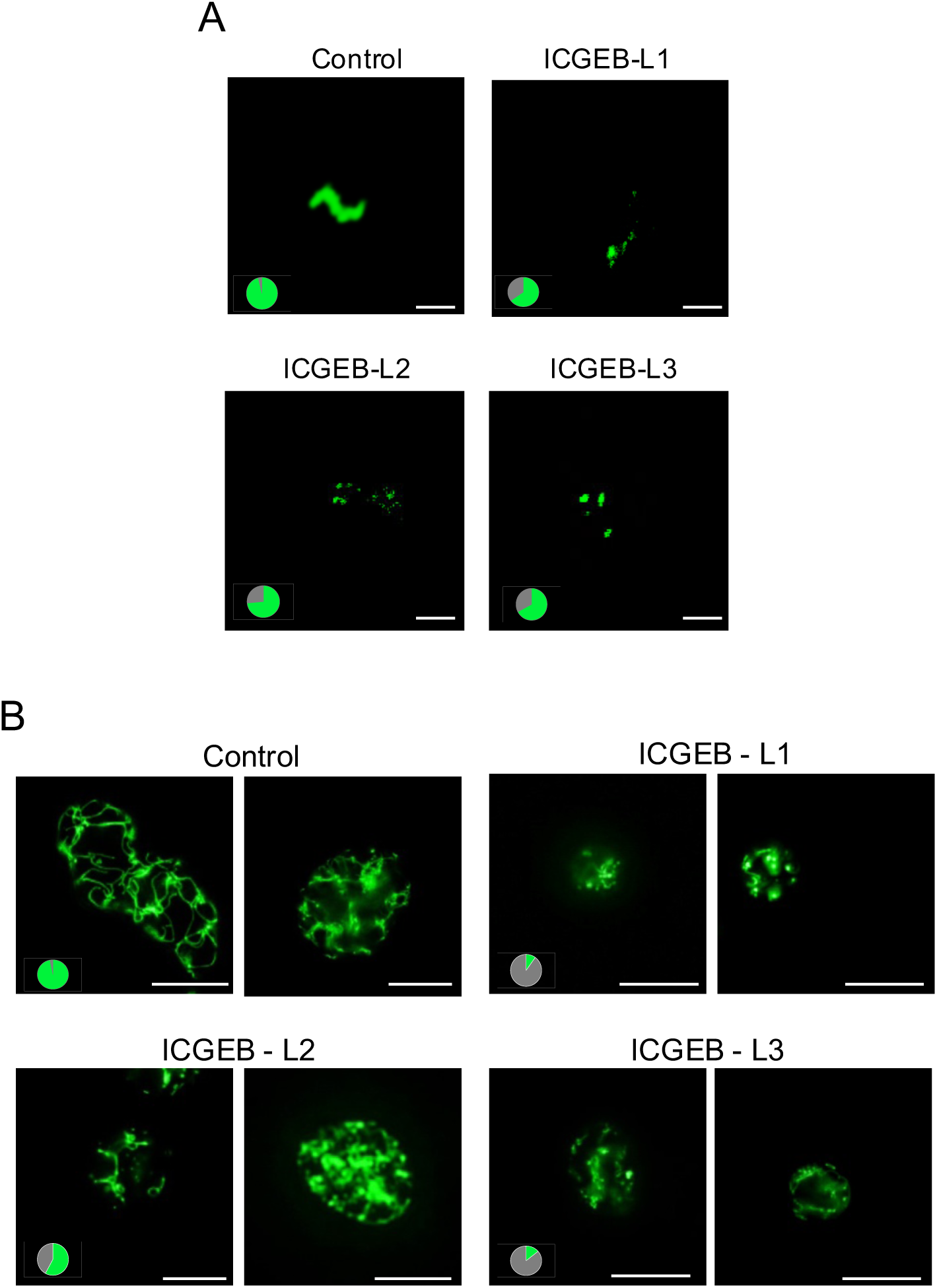
Selected lead compounds inhibit mitochondria development in the parasites. (A) Fluorescence microscopic pictures of GFP labelled mitochondria in P. falciparum blood stage parasites after treatment with each of the selected compound. (scale bar = 2μm) (B) Fluorescence microscopic pictures of GFP labelled mitochondria in *P. berghei* intraheptocytic stage parasites after treatment with the compounds. (scale bar = 10μm). Pie charts show proportion of parasites having normal (green) and abnormal/under-developed (grey) morphology of mitochondria in each set.

To assess the effect of compounds on mitochondrial development during intra-hepatocytic cycle, HepG2 cells were infected with *P. berghei* sporozoite (Pb-GFPmito, mitochondrial tagged parasite line; 14) and treated with each of the compound. As shown in Fig. 7C, the control sets showed normal development of the parasites mitochondria which showed highly branched network-like structures; whereas in treated sets, the parasites harboured underdeveloped or abnormal mitochondria. In ICGEB-L1 and ICGEB-L3 sets, >90% of the parasites showed abnormal or underdeveloped mitochondria (Fig. 7C, D).

### *In vivo* anti-parasitic efficacies for blood stage infection

The *in vivo* evaluation of efficacy for the test compounds was carried out using *P. berghei* mice malaria blood stage infection system using parasites expressing mCherry-luciferase, (Pb-mCherryhsp70Lucef1α;15). Infected mice were infused once daily by an intraperitoneal injection with compound or solvent alone for four consecutive days and parasite growth was analyzed. In the control set, the parasite growth was observed with a small peak on day 5 after infection, the parasitemia increased continuously from day 6 onward reaching 8-10% in all the mice by day 9 (Fig. 8A). Mice treated with ICGEB-L1 and ICGEB-L2 showed an initial increase in parasitemia, which was close to that of the control on day 5 and 6. However, growth of the parasites was suppressed from day 6 onwards and total parasitemia reduced to significantly low levels as compared to control. Mice treated with ICGEB-L3 showed much lower parasitemia on day 5 as compared to control; the parasites in these mice could not grow rapidly and these mice showed continuous low levels of parasitemia until day 9 in all experiments. Suppression of total parasitemia in treated mice as compared to control mice was calculated for day 5 and day 9. Mice treated with ICGEB-L1 showed ~50% and ~90% suppression of parasitemia on day 5 and 9 respectively, whereas mice treated with ICGEB-L3 showed ~80% and ~70% growth suppression on day 5 and 9 respectively (Fig. 8B, C). Overall the results show effective suppression of parasite growth in mice treated with ICGEB-L1 and ICGEB-L3 compounds.

**Figure 8:**
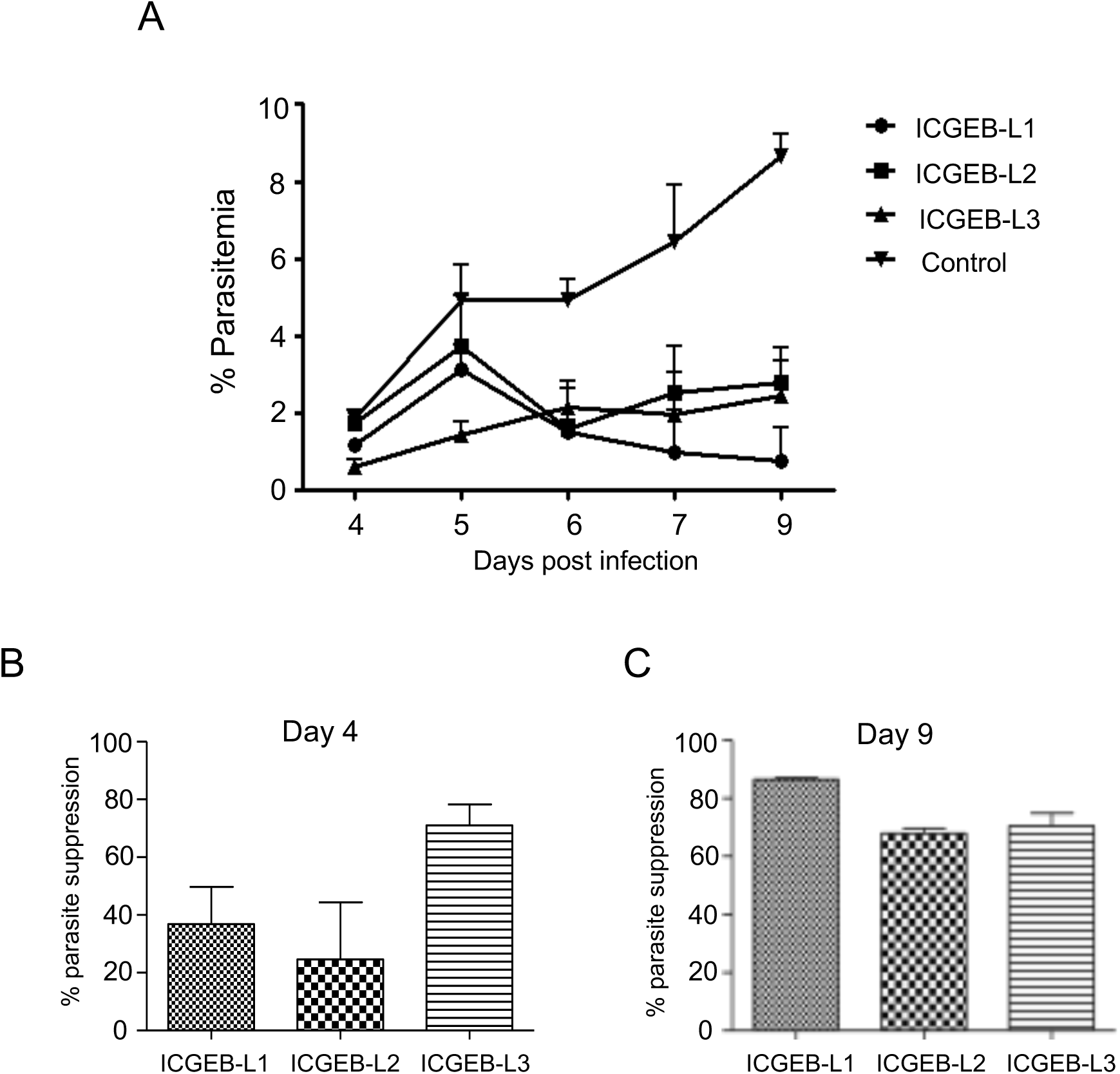
*In vivo* growth inhibition of blood stage parasite infection by treatment with selected compound in mice malaria model. (A) Graph showing parasitemia profile for average of five mice in the drug treatment set as compared to control set for each of selected compound. (B) Graph showing percentage parasite suppression (based upon mean parasitemia) at Day 4 and 9 as compared to control set for each of the drug treatment sets.

## Discussion

The metabolic pathways in the mitochondrion and the apicoplast, two parasite organelles of prokaryotic origin, are considered as suitable drug targets in the parasite. The mitochondrial ClpQ protease in *P. falciparum* is a homologue of prokaryotic ClpQ system, and thus it is considered as a potential drug target. We have earlier shown that the *P. falciparum* ClpQ, *Pf*ClpQ, harbours threonine protease activity; further the *Pf*ClpQ protease is localized in the parasite mitochondria (9, 10). Indeed, parasite proteases, especially organelle protease, are considered as important drug targets (16–21). Detailed studies by our group including a trans-dominant negative approach clearly showed that *Pf*ClpQ is essential for the parasite survival (9, 11); further these studies showed that the *Pf*ClpQ plays a key role in development of functional mitochondria and its segregation. The facts that the ClpQY is a prokaryotic machinery in the parasite having no homologue in human host, which is highly conserved across all the species of *Plasmodium*, combined with our data from genetic ablation studies, validated *Pf*ClpQ as a potent drug-target; this prompted us to identify *Pf*ClpQ specific inhibitors, which could be developed as new anti-malarial.

The Medicines for Malaria Venture (MMV, Switzerland) made unique initiatives to provide compound libraries, with proven biological activity, for drug discovery against neglected tropical diseases; one such initiative is the ‘Malaria Box’, which comprises of 200 probe-like compounds and 200 drug-like compounds having physiochemical properties in compliance with the Lipinski’s ‘rule of 5’. All the compounds of this library have proven anti-malarial activity on asexual blood stages of *P. falciparum*, together with a >10 fold selectivity ratio in the cytotoxicity assay on HEK293 cells (22). However, there is a need to identify the specific protein target of these compounds in the parasite, which his essentially required for SAR and medicinal chemistry campaign to further develop these compounds as lead anti-malarial. A number of studies have utilized the Malaria Box for activity assay based screening to identify hit and lead candidates targeting different enzymatic pathways in the parasite (23).

Here we have developed robust activity assay for the recombinant *Pf*ClpQ enzyme, that can be used for large scale screening of compounds. Screening of the ‘Malaria Box’ using this *in vitro Pf*ClpQ protease activity assay identified a potent inhibitor of *Pf*ClpQ with nanomolar efficacy, matching with its efficacy on the asexual blood stages. The hit compound is a hetercyclic compound having a core structure of a pyrimidotriazine. A number of studies have identified derivatives of pyrimidotriazine with varuous biological activities including anticancer, antimicrobial, antiviral and antifungal properties (24–26). In addition, there are reports of use of derivatives of pyrimidotriazine in treatment of asthma and diabetes (27, 28). Based upon the core structure of this hit compound, CN172, several derivatives were identified having variations in three of the side chains; primary screening of these derivatives generated SAR data of these derivatives based upon common scaffold.

Further screening of selected derivatives, which had better efficacy in inhibiting the enzyme as compared to hit compound, identified compounds having parasiticidal efficacies comparable to their efficacies on the enzyme. A number of compounds having better efficacy as compared to parent hit compound were not able to inhibit parasite growth, this could be due to poor solubility and uptake by the cells in the assays. Almost all the compounds which showed parasiticidal efficacies also showed high selectivity index on the mammalian cell lines. Overall these studies identified three potent active compounds (labelled as ICGEB-L1, ICGEB-L2 and ICGEB-L3 respectively) having 5-7 fold better efficacy than the parent hit compound, both in terms in enzyme inhibition and parasiticide effects, along with high selectivity index as compared to mammalian cells. The EC_50_ values of these compounds correlated with their respective IC_50_ on the enzyme activity. Further, all the three selected compound were found to be effective in mouse malaria model, showing effective suppression of parasitemia in 4-day suppressive test (29); specifically ICGEB-L3 showed ~75% suppression of parasitemia on day 4, whereas ICGEB-L1 and ICGEB-L2 showed >80% and ~60% parasite suppression on day 9 as compared to control. These results validate the selected compounds a potential candidate to develop further, through medicinal chemistry campaign, into new anti-malarials.

The Clp family of AAA proteases are shown to be regulatory proteases in prokaryotes and mitochondria (30–32). The ClpQ protease is localized in the matrix of the mitochondria in *P. falciparum* (10) as well as in *Trypanosoma brucei* (33). We have previously shown that the *P. falciparum* ClpQ protease (*Pf*ClpQ) is essential for parasite survival; trans-dominant negative effect of expression of active site mutant *Pf*ClpQ disrupted mitochondrial development in the parasite, which led to significant loss of mitochondrial membrane potential. These effects on mitochondrial growth and segregation were combined with ablation of parasite development at trophozoite/early schizont stages (9, 11). The morphological and growth phenotype in the blood stage parasites after treatment with selected compounds mimicked with that induced by the gene ablation studies of *Pf*ClpQ; which suggested specific targeting of *Pf*ClpQ in parasite mitochondria in the blood stage parasites.

Generation of drug resistance parasite has been reported for all commonly used antimalarials, including artemisinin. Therefore it is important to assess the propensity of parasite to generate resistance for a potential hit compound. Although the correlation between the rate of resistance generation in laboratory setting and that in the clinical setting is not established well, still the *in vitro* assay can help to understand the risk of generation of resistance under clinical studies. In our study the no resistant parasites were generated after growing in sequentially increasing concentration of each of the selected drugs. Only in one of the sets, the cloned parasite strain after first round of treatment (0.5×IC_50_), for ICGEB-L3 minor increase in the IC_50_ was detected. It has been shown a minimum of a 10-fold difference in IC_50_ between sensitive and resistant parasite is be a robust cut-off to exclude the false-positive resistance indication (34); therefore, these minor changes in IC_50_ observed in our studies are not significant. Overall, we show that the *P. falciparum* parasite show low propensity to develop resistance to selected hit compounds. However, all antimalarial therapies are developed as combination of two or more drugs, owing to general concerns about development of resistance in clinical settings as well as to improve the efficacies of the treatment. We also tested these compound in dose combination studies with known antimalarials; these studies identified potential partner drugs for combination therapies which can synergistically or additively increase the parasiticidal efficacies.

The hepatocytic cycle is the first stage of malaria infection in the human host; compound targeting key metabolic pathways required for rapid growth and segregation of liver stage parasite could have potential use as prophylactics. Efficient and accurate process of cytokinesis and organelle development and segregation is crucial for liver stage parasites; during the liver stage, the parasite undergoes extensive replication, forming tens of thousands of infectious merozoites in each infected hepatocyte. To cope with extensive segregation the parasite organelles need to be highly metabolically active, divide and segregate extensively; indeed, it has been shown that during the growth phase of liver stage parasites, the apicoplast and mitochondrion become two extensively branched and intertwining structures in parallel with nuclear division (14). *Pf*ClpQ is the only protease machinery shown to be localized in the mitochondria matrix which gets expressed in all the parasite stages, therefore we also assessed anti-parasitic potential of the *Pf*ClpQ inhibitors on the hepatocytic stages of the parasites. All the three selected hit compounds were found to be effective on growth and development of liver stages, and their potency was much higher than that on the blood stages, which could be due to extensive metabolic activity of the mitochondria in the liver stages. Therefore, we show that the *Pf*ClpQ targeting compounds hold promise as prophylactic and chemoprotective agents for malaria.

In summary, the MMV Malaria Box was subjected to robust activity-based screening to identify hit compound targeting essential prokaryotic protease in *Plasmodium*. The SAR analysis identified potential hit compounds with effective parasiticidal efficacies against both blood and hepatocytic stages of the parasite. Initial *in vivo* studies confirm these inhibitors as potential antimalarial drug candidates. These compounds can be starting point for medicinal chemistry campaign, and the SAR data provides essential information required for further drug development.

## Materials and Methods

### Expression and purification of recombinant protein

The recombinant ClpQ protease was expressed and purified as described earlier (35, 36). Briefly, a gene fragment corresponding to mature region of ClpQ protein (112-621 bp; 38-207 aa) was cloned into pET29a vector (Novagen) to obtain the plasmid construct pET29a-PfClpQ, which was transformed into *E. coli* expression cells BL21 (DE3) for expression of recombinant protein with C-terminal histidine tag. The recombinant protein was purified from bacterial cell lysate by affinity chromatography using Ni-nitrilotriacetic-acid (Ni-NTA) agarose resin (Qiagen). Eluted fractions were subjected to SDS-PAGE and western blot analysis to assess the purity of the purified recombinant protein.

### Protease activity assay and Enzyme kinetics

*In vitro* fluorometric assays for the threonine protease activities was carried out using recombinant proteins and specific fluorogenic peptide substrates, Suc-Gly-Gly-Leu-AMC; the activity was determined by estimating release of 7-amido-4-methylcoumarin (AMC) from the peptide substrate as the increase of fluorescence (excitation 355 nm; emission 460 nm) using a Victor-3 Fluorometer (Perkin-Elmer). Initial standardization of the assay was carried out using different assay buffers at a pH range from 5-9 with varying concentrations of the reductant DTT (0-10mM). The assay conditions with maximal protease activity were used; the assays were carried out in 200 µl reaction volume in a 96 well plate format (NUNC) containing 0.8µM of recombinant protein. The fluorogenic peptide substrate (Suc-GGL-AMC) was added at 90 µM final concentration and the release of AMC was continuously monitored for 3-6hr at room temperature. Threonine-protease inhibitor N-(benzyloxycarbonyl)-leucinyl-leucinyl-leucinal (MG132) containing wells were also set up under similar assay conditions to confirm the specificity of the reaction.

To assess robustness of the activity assay, HML test was conducted for a period of three days in accordance for statistical test to calculate the Z-value. The HML assay employed three different reaction wells: the **<u>H</u>**igh activity well i.e. protease activity well without any inhibitor; **<u>M</u>**edium activity well i.e. in the presence of inhibitor, MG132 (14µM final concentration); and the **<u>L</u>**ow activity well that is the substrate control well without the protein; this set of three wells (HML) was repeated throughout the 96-well plate. The assay was repeated for three consecutive days. The Z-value was calculated based on the change in fluorescence after 6 hrs.

For calculating the kinetic parameters *K*m and *V*max varied concentration of the peptide substrates at constant enzyme concentration (0.8µM) was set up in a reaction plate in duplicates. Rate of peptide hydrolysis was recorded for 6 hrs at every one-hour interval. The kinetic constant *K*m and *V*max were determined by fitting Michalies-Menten curve using the Graph Pad Prism V5.0 software package. The inhibitory efficacies of compounds were assessed in the same assay, carried out in presence or absence of the chemical compound at different concentration (0-1000nM). The IC50 values for each of the compound were calculated from curve fittings by software Workout V 2.5.

### Compound library screening, identification of hit and lead compounds

The ‘Malaria Box’ 400 compounds consists of 200 drug-like and 200 probe-like anti-malarial compounds. These 400 compounds were selected from a list of more than 20,000 compounds (22); further, cytotoxicity of these compound was assessed on human embryonic kidney cells (HEK 293), with reference to the controls treated with puromycin. The full list of the 20,000 hits from *P. falciparum* whole cell screening originates from the GlaxoSmithKline Tres Cantos Antimalarial Set (TCAMS), Novartis-GNF Malaria Box Data set and St. Jude Children’s Research Hospital’s Dataset available on the ChEMBL-NTD website (22). The ‘Malaria Box’ compound library was obtained from MMV, Switzerland. The ‘Malaria Box’ compounds as well as the derivatives of hit compound identified were screened using the standardized *in-vitro* protease activity assay for assessing their potential as *Pf*ClpQ inhibitors. Briefly, the recombinant enzyme (5µg) was incubated in the assay buffer with different concentrations of each of the compound or DMSO alone; the reactions were initiated by addition of the activity assay reaction mixture to a final volume of 200µl and the substrate hydrolysis was monitored as described above.

### *P. falciparum* parasite culture, growth inhibition analysis

*P. falciparum* was cultured in RPMI media (Invitrogen) supplemented with 0.5% (w/v) Albumax™ (Invitrogen) 4% haematocrit. Culture was kept static in a gas mixture of 5% carbon dioxide, 5% oxygen and 90% nitrogen at 37°C. To assess the effect of selected inhibitors on parasite growth, Synchronous culture with ring stage parasites (6-8 hpi) at ~1% parasitemia was grown in 24-well flat-bottomed cell culture plates. The compound was added to the parasite cultures at the desired final concentrations (0-1000 nM). A control well was also set up with solvent only as negative control. Each growth inhibition assay was performed in triplicate, and the experiment was repeated twice. After 48h, growth of the parasite was assessed by DNA fluorescent dye-binding assay using SYBR green (Sigma) following (37). The effect of these compounds on growth of the mammalian cell lines was assessed using standard cell proliferation assays as described earlier (36).

### Fluorescence microscopy to assess mitochondria development

To assess the effect of selected compounds on growth and development the parasite mitochondria in blood stages, we utilized *P. falciparum* parasites expressing HSP60-GFP fusion protein (13). Tightly synchronized parasites at ring stages (6-8hpi) were treated with each of the compound and after 24h of growth (trophozoite stages) these parasites were stained with 40,6–diamidino-2-phenylindole (DAPI, Sigma) at a final concentration of 2 μg ml-1 for 10 min, and visualized under the confocal microscope.

To assess the effect of compounds on mitochondrial development during intra-hepatocytic cycle, HepG2 cells were infected with sporozoite of mitochondrial tagged parasite line *P. berghei* (Pb-GFPmito) (14) were used; the infected cells were treated with each of the compound. The GFP labelled mitochondria in asexual and intra-hepatocytic parasites were viewed using a Nikon A1 Confocal microscope, images were acquired with Plan Apochromat 100×/1.40 NA oil immersion objective lens and analysed by NiKDn-nis element software (version 4.1).

### *Ex vivo* anti-parasitic efficacies of compounds against *P. berghei* parasites

To study the in vivo anti-parasitic efficacies of compounds, we first assessed their effect on growth of *P. berghei* blood stage parasites *in vitro* using a schizont development assays following (38). Briefly, ring stage *P. berghei* parasites were grown overnight in RPMI media with 10% FBS to develop into mature trophozoite/schizont stages. Each of the compounds was added to the media in varying concentrations (10-300nM) and percentage schizont development was compared to that in control set.

### *In vivo* anti-parasitic efficacies of selected compounds on blood stage infection

The *in vivo* evaluation of efficacy for the test compounds was carried out using *P. berghei* mice malaria blood stage infection system. Briefly, 6-to 8-week-old female Balb/C mice were infected with *P. berghei* parasites expressing mCherry-luciferase, (Pb-mCherry_hsp70_FL_ef1α_) (15). Mice were monitored daily for the presence of parasites by examining blood sample under the fluorescence microscope for mCherry-expressing parasites. Once the presence of parasites was confirmed, mice were infused once daily by an intraperitoneal injection with compound (15 mg/Kg body weight for ICGEB-L1; 60 mg/Kg body weight for ICGEB-L2; and 40 mg/Kg body weight for ICGEB-L3) or solvent alone for four consecutive days. Parasitemia was determined by light microscopy, by examining a Giemsa-stained peripheral blood film for each mouse. In addition, parasite growth was also monitored by *in vitro* luciferase assays using infected mice blood samples collected at different time-points and Dual-Luciferase Reported Assay System (Promega). During the experiments any animal that was judged to be ill were promptly euthanized.

#### Intra-hepatocyte parasite growth and development assay

The compounds were assessed for their anti-parasitic efficacies on intra-hepatocytic stages using *in vitro P. berghei* sporozoite (Pb-mCherry_*hsp70*_, parasites constitutively expressing mCherry in the cytosol for detection, (12) infection of HepG2 cells. Briefly, 5×10^4^ HepG2 cells per well were seeded in 96 well plates to generate a confluent cell layer. The next day, *P. berghei* sporozoites isolated from salivary glands of infected *Anopheles stephensi* mosquitos were added to the cells; 2h after infection the cells were detached using the mild detachment enzyme Accutase. 1:8 dilutions of the cells were seeded into wells containing different drug concentrations (the drug solvent DMSO was used as control). The development of exoerythrocytic forms (EEFs) was analysed using the automated imaging system ‘IN Cell Analyzer 2000’ (GE healthcare) by counting the parasite number and measuring parasite size at 24 and 48 hpi as described earlier; infected cells in each well (*n*=8) were analyzed and mean parasite size was determined for each set. The potency of each drug to inhibit parasite EEF growth was compared to the control set. The effect of each drug was assessed at the IC_50_ concentration determined for that drug for *P. falciparum* blood stages (Pf-IC_50_), 3× lower and 3× higher. The assay was conducted twice, with similar outcome in both experiments.

The effect of different treatments (each compound at concentration of Pf-IC_50_, 3× lower and 10× lower) was examined with the output being completion of liver stage development, as determined by detached cell formation at 65hpi. as described earlier (12). Briefly, the assays were set up as described above, at 65hpi the supernatant was collected from infected well and the number of detached cells in the supernatant was counted using a fluorescence microscope in triplicate. To study the effect of the compounds on different developmental stages of intra-hepatocyte parasites, the compounds were added for varying time periods after sporozoite infection of HepG2 cells (schematic representation Figure 6A) with the output also being completion of liver stage development (detached cell analysis).

### Statistical Analysis

The data sets were analysed using GraphPad Prism ver 5.0 to calculate *K*m, *V*max, IC_50_ and EC_50_ values, and the data were compared using unpaired Student’s *t*-test.

## Declarations

### Funding

This study was supported by “Malaria Box Challenge Grant” (MMV 14/6008/02) by Medicines for Malaria Venture (MMV, Switzerland) to AM and VH. The research work in AM laboratory is supported by Flagship Grant (#BT/IC-06/003/91) and by Centre of Excellence grant (BT/COE/34/SP15138/2015) from the Department of Biotechnology, Govt. of India.

## Acknowledgements

The “Malaria Box” compound library was provided by Medicines for Malaria Venture (MMV, Switzerland). We are grateful to Siggi Sato for providing plasmid constructs (pSSF2-HSP60-GFP) for targeting GFP to the malarial mitochondrion. We thank Rotary blood bank, New Delhi, for providing the RBCs.

## Animal Ethics approval

The research study was approved by the Institutional Animal Ethics Committees of ICGEB-New Delhi and ICB, Bern.

## Notes

### Competing Interest Statement

The authors have declared no competing interest.

